# Metallothionein-3 attenuates the effect of Cu^2+^ ions on actin filaments

**DOI:** 10.1101/2022.09.23.509211

**Authors:** Rabina Lakha, Carla Hachicho, Matthew R. Mehlenbacher, Dean E. Wilcox, Rachel N. Austin, Christina L. Vizcarra

## Abstract

Metallothionein 3 (MT-3) is a cysteine-rich metal-binding protein that is expressed in the mammalian central nervous system and kidney. Various reports have posited a role for MT-3 in regulating the actin cytoskeleton by promoting the assembly of actin filaments. We generated purified, recombinant mouse MT-3 of known metal compositions, either with zinc (Zn), lead (Pb), or copper/zinc (Cu/Zn) bound. None of these forms of MT-3 accelerated actin filament polymerization *in vitro*, either with or without the actin binding protein profilin. Furthermore, using a co-sedimentation assay, we did not observe Zn-bound MT-3 in complex with actin filaments. Cu^2+^ ions on their own induced rapid actin polymerization, an effect that we attribute to filament fragmentation. This effect of Cu^2+^ is reversed by adding either EGTA or Zn-bound MT-3, indicating that either molecule can chelate Cu^2+^ from actin. Altogether, our data indicate that recombinant MT-3 does not directly bind actin but it does attenuate the Cu-induced fragmentation of actin filaments.

## Introduction

Mammalian metallothioneins (MTs) are small (6-8 kDa), cysteine-rich proteins that bind seven *d^10^* metal ions in two metal-thiolate clusters (Figure 1; [1]). Their physiological roles include metal detoxification and metal homeostasis, and toward this function MTs interact with both essential (Zn^2+^, Cu^+^) and abiotic (Hg^2+^ and Cd^2+^) metals (reviewed in [2,3]). Among the four mammalian MT isoforms, metallothionein-3 (MT-3) has a distinct expression pattern: it is found predominantly in the central nervous system [4,5] and kidney [6]. Unlike most MTs [7], MT-3 expression is not induced by zinc [8,9] but is upregulated by hypoxia [10–12]. While MT-3 deletion in the mouse has little apparent effect on overall health and cognition, *Mt3*-null mice do have an enhanced sensitivity to seizures induced by kainate, a neuroexcitatory glutamate analog [13]. Compared to wild type mice, *Mt3*-null mice have similar levels of copper in the brain but less zinc in certain brain regions, including the hippocampus [13]. To our knowledge, kidney function and metal loads have not been studied in the *Mt3*-null mice.

**Figure 1.**
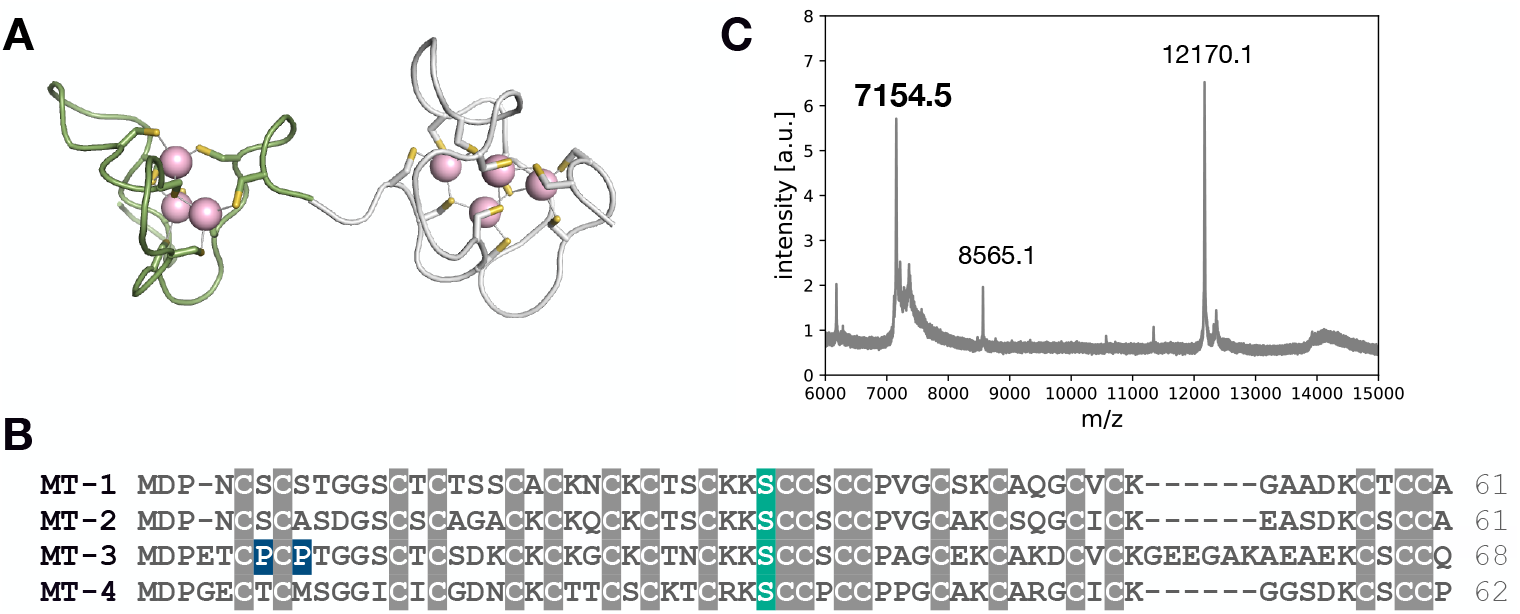
Structure and sequence alignment of metallothioneins (MTs). (A) The solution structure of MT-2 is shown with cadmium bound. The N-terminal β-domain (green; 2MHU) and the C-terminal α-domain (gray;1MHU) have three and four Cd^2+^ ions bound, respectively. (B) The four mouse metallothioneins were aligned using accession numbers: NP_038630.1 (MT-1), NP_032656.1 (MT-2), NP_038631.1 (MT-3) and NP_032657.1 (MT-4). Cysteine residues are highlighted in gray. The two mutated prolines in the non-GIF P7S/P9A mutant are highlighted in blue, and the serine mutated in the S33D mutant is highlighted in green. The numbers on the right of the sequence indicate the number of amino acids in each MT. (C) MALDI of the purified MT-3 showing the molecular weight at 7154.5 Da. The predicted molecular weight is 7153.3. The other two masses are from calibration buffer.

MT-3 has a unique biological activity: it inhibits brain extracts from promoting the survival of cultured neurons [14], an activity that is not shared by other MT isoforms [15,16]. In fact, the *Mt3* gene product was initially named growth inhibitory factor (GIF) before it was renamed when molecular cloning revealed its homology to canonical MTs [4,17]. While brain extracts from both Alzheimer’s disease (AD) patients and aged-matched controls are neurotrophic, the AD brain extract is approximately two-fold more potent than control extracts at promoting survival of cultured neurons [14,15]. Some have attributed this difference to lower levels of MT-3 in AD brain extracts [17,18], while others have found that MT-3 levels are unchanged in AD [15,19]. A molecular mechanism of growth inhibition by MT-3 has not been elucidated. However, the domain and MT-3-specific sequence motifs that are essential for GIF activity have been mapped. MT-3’s N-terminal *β* domain is necessary for growth inhibition [16,20]. The C-terminal *α* domain may be dispensable for GIF activity [16], or it may stabilize the *β* domain by domain-domain interactions [21]. Mutational studies identified two conserved prolines (Pro7 and Pro9) and a threonine (Thr5) that are unique to MT-3 as essential for GIF activity [16,22,23]. Since the growth inhibition bioassay reflects MT-3’s attenuation of brain extract-induced growth stimulation [15,16], it is likely that MT-3 interacts specifically with some component of the extract or with a neuronal receptor.

Motivated by the potential importance of protein-protein interactions in MT-3 function, several screens have been carried out to characterize MT-3’s interactome [24–26]. Using immunoaffinity chromatography and tandem mass spectrometry, at least 12 proteins have been identified that bind to MT-3 [24,25]. Of particular interest to this work, the cytoskeletal protein actin (*β* and *γ* isoforms) was one of the proteins most often identified in complex with MT-3 in these screening studies [24–26]. Based on the list of MT-3-binding proteins, Armitage and coworkers proposed that MT-3 is secreted from astrocytes through its interaction with Rab3A, Exo84p, and 14-3-3 zeta and then transported into neurons through interaction with plasminogen (PGn) and enolase, where MT-3 could inhibit growth through its effect on the cytoskeleton and/or other interactions [25].

In addition to the appearance of actin in interaction screens, MT-3 and actin have been functionally linked in mouse astrocytes [27,28] and in human kidney proximal tubule cells [26]. In astrocytes, changes in the actin cytoskeleton that occur downstream of epidermal growth factor (EGF) treatment are abnormal in the absence of MT-3 [27]. In the same study, two observations supported a direct binding interaction between MT-3 and actin. First, GFP-MT-3 was detected in the pellet fraction after high-speed centrifugation of astrocyte lysates, consistent with an interaction between GFP-MT-3 and actin filaments (‘F-actin’). Second, GFP-MT-3 isolated from astrocytes enhanced actin polymerization in an *in vitro* pyrene-actin polymerization assay [27]. Addition of the zinc chelator TPEN blocked MT-3-induced actin polymerization, while the addition of Zn-bound TPEN did not, suggesting that MT-3 must be in a Zn-bound and not a Cu-bound state to accelerate actin assembly [27].

The connection between Zn and MT-3-induced actin polymerization [27] is surprising because the *β* domain of MT-3 isolated from brain is Cu-bound [29], giving an overall metal stoichiomentry of Cu_4_Zn_3,4_MT-3 [17]. Furthermore, MT-3 is the most copper-philic of the mammalian MTs [30,31]. In general, copper is an essential and highly regulated trace element with a variety of roles in normal growth, development, and cellular metabolism [32] and is known to bind to both actin [33] and MT-3 [31,34]. In hippocampal neurons, copper is enriched in dendrites compared to somata, where nano X-ray fluorescence suggests it co-localizes with actin filaments in dendritic spine necks [35]. Both copper supplementation with added CuSO_4_ and copper chelation by neocuproine resulted in decreased actin filament content in dendrites [35], suggesting that Cu levels impact the actin cytoskeleton. *In vitro*, it was found that Cu^2+^ binds with subpicomolar affinity to monomeric actin through Cys374, an interaction that still allows actin to polymerize into filaments [33]. Electron microscopy studies showed that Cu^2+^ ions induce fragmentation of actin filaments, leading to shorter filaments but not complete depolymerization [36]. The Cu-binding properties of MT-3 have been linked to its neuronal protective abilities: MT-3 can swap its bound Zn^2+^ for Cu^2+^ that is bound to neurologically important peptides like amyloid-β peptides, a-synuclein, and prion protein, thereby eliminating harmful redox chemistry associated with copper [37–40].

Despite evidence for an interaction between actin and MT-3, it is unclear whether they bind directly or their interaction is mediated by other proteins, metal ions, and/or post-translational modification. If actin and MT-3 do form a stable protein-protein complex, it is unknown whether this is critical to MT3’s GIF activity and potential neuro-protective roles. MT-3’s role as a metal chelator and metal chaperone suggest other mechanisms by which MT-3 and actin might impact each other’s functions. Careful biochemical characterization of the MT-3-actin interaction is required to better understand the role of MT-3 in the central nervous system. Such characterization must consider the different potential metalation states of MT-3, as well as the possibility of *in vivo* phosphorylation in mammalian systems. In this study, we show that while recombinantly-expressed MT-3 has minimal direct effect on actin filament dynamics, MT-3 modulates the effect of Cu^2+^ on actin filaments.

## Methods

Mouse MT-3 (UniProt P28184) was expressed in BL21-DE3-pLysS *E. coli* cells and purified as previously described [34]. To produce different metalation states (Zn_7_MT-3, Pb_7_MT-3, Cu_4_Zn_4_MT-3), stock solutions of either ZnCl_2_, Pb(NO_3_)_2_, and/or CuCl_2_ were added to apo-MT-3 in stoichiometric amounts. A non-GIF mutant (Pro7Ser/Pro9Ala) and a phospho-mimetic mutant (Ser33Asp) were both constructed by PCR-based mutagenesis [41] (DNA sequences in SI) and purified using the same protocol as WT MT-3 [34]. Rabbit skeletal muscle actin from acetone powder (*Pel-Freez Biologicals*) and recombinant *S. pombe* profilin were purified, and actin was labeled as previously described [42].

### Fascin purification

Fascin protein was expressed in BL21-DE3* *E. coli* cells. Terrific broth (1 L supplemented with 100 μg/mL ampicillin) was inoculated with 10 mL of dense culture grown for 16 hr at 37 °C. These large cultures were shaken at 250 rpm and 37 °C until the cells reached the optical density of 0.3–0.6 at 600 nm. The cultures were then chilled to 18 °C for 30 min, and protein expression was induced with 0.25 mM isopropyl *β*-D-1-thiogalactopyranoside (IPTG). The cells were harvested by centrifugation, resuspended in PBS (140 mM NaCl, 2.7 mM KCl, 10 mM Na_2_HPO_4_, and 1.8 mM KH_2_PO_4_ pH 7.3), flash frozen in liquid nitrogen, and stored at −80°C. To isolate fascin protein, 20 g of frozen cells were thawed and brought to 30 mL with lysis buffer (50 mM Tris-HCl, 500 mM NaCl, pH 8.0) supplemented with 1 mM phenylmethylsulfonyl fluoride (PMSF), 1 mM dithiothreitol (DTT) and 4 μg/mL DNase (*Sigma catalog no. DN25*). Cells were lysed using a French press and the lysate was centrifuged at 30,000 × *g* for 30 min. The supernatant was nutated with 2 mL glutathione Sepharose resin slurry (*Cytiva*) for 1 hr. The slurry was poured into a column and the resin was washed first with the lysis buffer followed by a low salt buffer (50 mM Tris-HCl, 100 mM NaCl, 1 mM DTT, pH 8.0). Protein was then eluted with glutathione elution buffer (50 mM Tris-HCl, 100 mM NaCl, 100 mM glutathione, 1 mM DTT, pH 8.0) and dialyzed into low salt buffer at 4 °C overnight. His_6_-tagged TEV protease (purified as described [34]) was added to the protein in dialysis tubing to cleave the fusion protein. The sample was nutated with glutathione Sepharose for 1 hr to separate the cleaved fascin from GST. The unbound protein was collected and nutated with Ni-NTA resin for 1 hr. The unbound fraction was collected from the Ni-NTA column and dialyzed into KMEI dialysis buffer (50 mM KCl, 1 mM EGTA, 1 mM MgCl_2_, 10 mM imidazole, 1 mM DTT, pH 7.0) overnight. The protein was concentrated using the Amicon 30 kDa centrifugal filter, and the final concentration was measured using an extinction coefficient of 68,465 M^−1^cm^−1^ at 280 nm. Proteins were aliquoted in small fractions, flash-frozen in liquid nitrogen, and stored at −80 °C.

### MALDI-TOF mass spectrometry

Matrix-assisted laser desorption ionization time of flight (MALDI-TOF) mass spectrometry was carried out using a Bruker ultrafleXtreme MALDI TOF/TOF instrument fitted with a frequency-tripled Nd:YAG laser (355 nm). The matrix contained 10 mg/mL sinapinic acid in 70:30 water:acetonitrile with 0.1% trifluoroacetic acid. Each protein sample was prepared in the ratio of 3:7 protein:matrix, and 1 μL of that mixture was spotted onto a stainless steel sample plate followed by drying at room temperature. The data were collected in linear positive-ion mode.

### Actin co-sedimentation and monobromobimane labeling

Actin co-sedimentation assays were performed to test for an interaction between Zn_7_MT-3 and F-actin. Monomeric actin was first polymerized by adding KMH polymerization buffer (1× concentrations: 50 mM KCl, 1 mM MgCl_2_, 10 mM HEPES, pH 7) and incubating at room temperature for 30 min. The F-actin was then incubated with either a range of Zn_7_MT-3 concentrations (20, 10, 5 and 2.5 μM) or fascin (5 μM) for 1 h at room temperature. The samples were then centrifuged at 100,000 ×*g* at 4 °C for 90 min. The supernatants were removed, and the pellets were resuspended in monobromobimane (mBrB) labeling buffer (60 mM Tris-Cl, 20 mM EDTA, 20 mM TCEP, pH 8.0) [43]. Equal volumes of mBrB labeling buffer and supernatant were mixed. A stock solution of 100 mM mBrB dye in acetonitrile was added to each sample to reach a final mBrB concentration of 12.5 mM. The samples were then incubated at room temperature in the dark for 1 hr. Excess mBrB was removed by extraction into methylene chloride. The samples were analyzed by SDS-PAGE (*Bio-Rad* AnyKD gel, 160V, 45 min) and Coomassie stained.

### Actin assembly assays

Bulk actin assembly assays were carried out as described [42], with a few changes. EGTA was excluded from the polymerization assays to avoid metal chelation. The polymerization buffer used was KMH (50 mM KCl, 1 mM MgCl_2_, 10 mM HEPES, pH 7.0) with 0.2 mM ATP. For the assays shown in Fig. S1, 10 mM MOPS was used in place of HEPES. In specific cases, EGTA, CuCl_2_, and/or ZnCl_2_ was added to the assay as indicated. Fluorescence was monitored using a TECAN F200 plate reader with an excitation filter at 360 nm (35 nm bandwidth) and an emission filter of 415 nm (20 nm bandwidth). To collect the images in Fig. 2C, ≈10 μL of reaction mixture was removed from the pyrene-actin polymerization assay at 50 s and 1500 s. These samples were mixed with Acti-stain488 phalloidin (*Cytoskeleton, Inc.*) at an equimolar actin to phalloidin ratio, diluted 50-fold into imaging buffer (1× KMH, 5 mM DTT), and immobilized on poly-L-lysine coated coverslips. Imaging conditions are described below.

**Figure 2.**
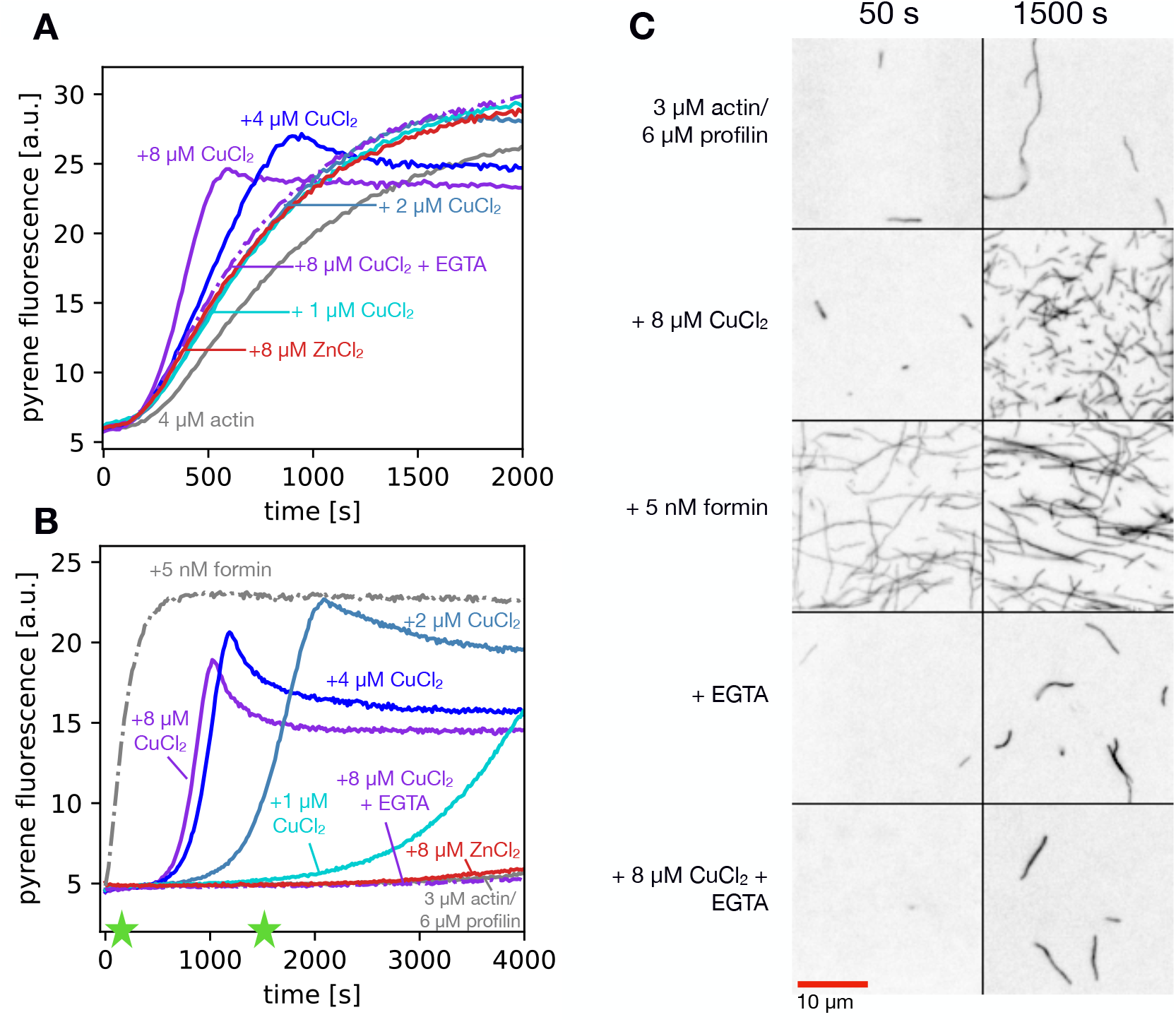
Effects of copper on actin polymerization. Pyrene-actin polymerization was measured in the presence of CuCl_2_, ZnCl_2_ and/or EGTA with (A) 4 μM actin or (B) 3 μM actin and 6 μM profilin. In all experiments, 5% of actin monomers were pyrene-labeled. In the presence of copper ions, a rapid increase in pyrene signal was observed after an initial lag period. This effect was particularly pronounced in the presence of profilin. The chelator EGTA (1 mM) reversed the effect of CuCl_2_ on actin polymerization. (C) Actin filaments were imaged using TIRF microscopy, sampled at two time points in the profilin/actin polymerization reaction. The green stars indicate the times (50 s and 1500 s) when a small amount of actin was removed from the assay and mixed with equimolar Acti-stain488 phalloidin, diluted 50× into imaging buffer, and immobilized on poly-L-lysine coated coverglass.

### ICP mass spectrometry

After the co-sedimentation assay, separated supernatants and pellets were mixed with equal volumes of 68% trace metal grade HNO_3_ overnight. The digested samples were diluted 400-fold by adding 10 mL of milliQ water. Samples were analyzed on a Multi-element ICP-MS (Element XR). The mass percentage of Cu in the pellet was calculated using the ppb of the pellet fraction, obtained from ICP-MS, divided by the total ppb of the pellet and the supernatant, multiplied by 100.

### Fluorescence microscopy

Monomeric actin was polymerized to form F-actin for 30 min at room temperature in 1× KMH. Samples were prepared by mixing 5 μM F-actin with 5 μM Cu^2+^ as indicated, followed by addition of 5 μM EGTA or 5 μM Zn_7_MT-3. Each sample was labeled with 2:1 Acti-stain488 phalloidin:actin, and further diluted by adding 1900 μL of imaging buffer to reach the final concentration of 6.5 nM F-actin and 12.5 nM phalloidin. Slides were prepared from each sample by pipetting 10 μL of sample onto a poly-L-lysine-coated coverslip. Samples were visualized using a Nikon Ti2 inverted microscope equipped with a TI-LA-HTIRF module and an ORCA-Flash 4.0 LT+ sCMOS camera, controlled with NIS-Elements software. An apochromat TIRF 100 × oil-immersion objective lens (N.A. 1.49) was used with a 400 ms exposure time and 2% (Fig. 2) or 5% (Fig. 3) power with a 20-mW 488 nm argon laser. All image analysis was conducted using the FIJI distribution of the ImageJ software [44].

### Isothermal titration calorimetry

Zn_7_MT-3 samples in 100 mM BisTris, 150 mM NaCl, pH 7.4 were stored and sample preparation occurred within a Coy Laboratory glovebox that maintains a N_2_ environment, with daily addition of H_2_. A platinum catalyst, which reacts with trace O_2_, and the H_2_, to form water, maintains low ppm oxygen. All buffers were made metal-free by soaking with Chelex resin (*Sigma*), filtered, and thoroughly degassed under vacuum before transfer into the glovebox. Diethylenetriaminepentaacetic acid (DTPA) was weighed into a borosilicate glass vial and brought into the glovebox for dissolution in deoxygenated buffer. The DTPA concentration was confirmed with ITC by titration with a known concentration of Zn^2+^. The experimental stoichiometry and binding enthalpy were compared to known values.

**Figure 3.**
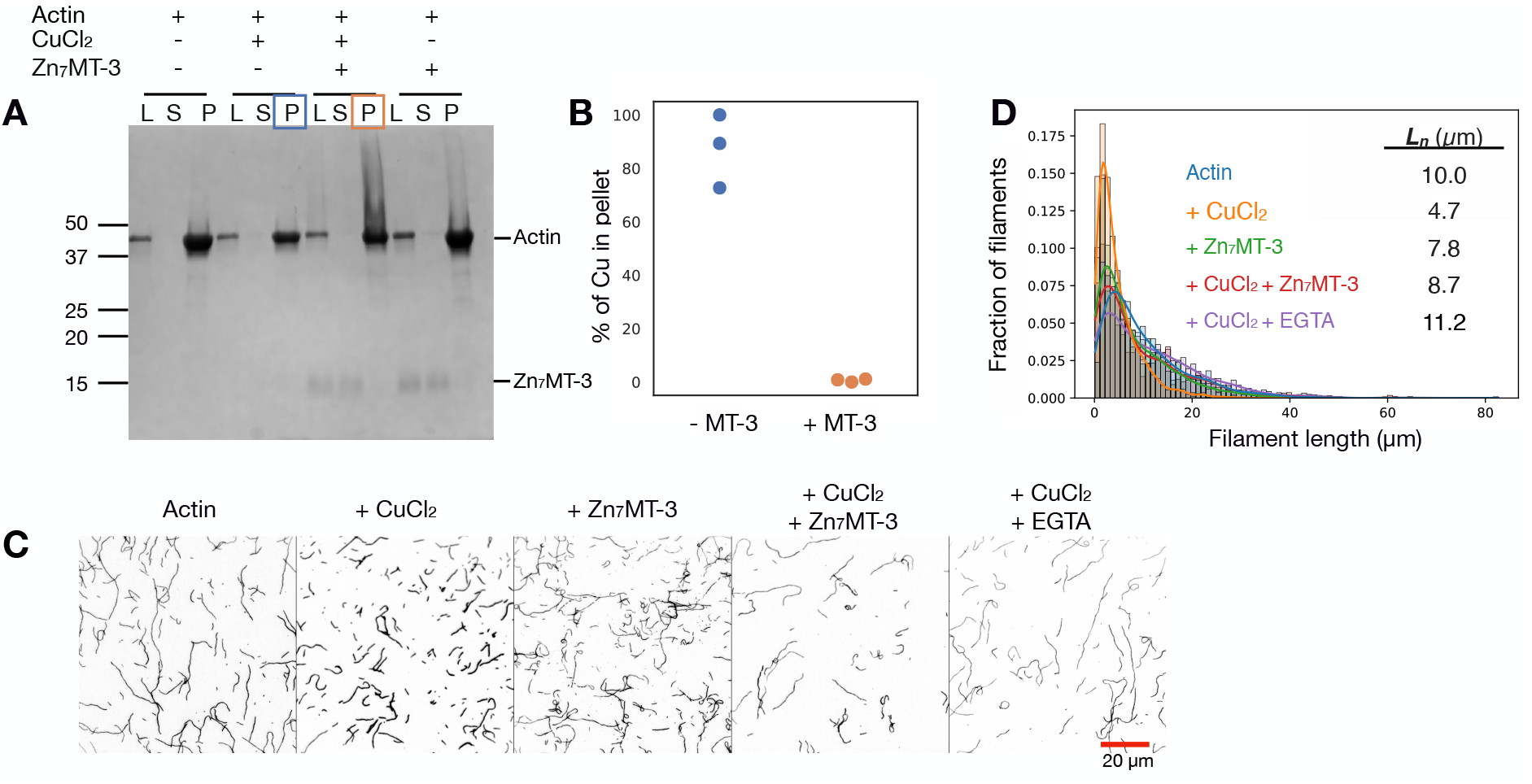
Copper ions induce actin filament fragmentation and MT-3 reverses it. (A) Co-sedimentation assay of 5 μM F-actin with CuCl_2_ (5 μM) and/or Zn_7_MT-3 (5 μM). After separating supernatant (S) and pellet (P) fractions, samples were labeled with monobromobimane, separated by SDS-PAGE, and stained with Coomassie. Under these conditions, there is no detectable Zn_7_MT-3 in the pellet. The pellets were concentrated 4-fold compared to the load (L) and supernatants. (B) ICP-MS quantification of the amount of copper ions in pellet (P) in the presence and absence of Zn_7_MT-3. Pellets boxed with blue and orange in panel A were analyzed in panel B. (C) Actin filaments were imaged using TIRF microscopy in the presence of different combinations of CuCl_2_, Zn_7_MT-3, and EGTA. All samples were mixed with Acti-stain488 phalloidin, before 50-fold dilution into imaging buffer and immobilization on poly-L-lysine coated coverglass. Actin filaments appeared fragmented when exposed to CuCl_2_. This effect was reversed with the addition of Zn_7_MT-3 or EGTA. (D) Histogram of filament lengths for each condition along with the average length (*L_n_*). The numbers of filaments counted for each condition were 1175 (Actin only), 1626 (+ CuCl_2_), 808 (+ Zn_7_MT-3), 541 (+ CuCl_2_ + Zn_7_MT-3), and 897 (+ CuCl_2_ + EGTA).

All ITC experiments were completed with a Malvern Panalytical (MicroCal) VP-ITC, which is housed in a custom-built glovebox that is purged with N_2_ to ensure an anaerobic environment. The ITC measurements used injection volumes of 4 or 5 μL, a stir speed of 307 rpm and a temperature set at 25 ± 0.2 °C. The heat of the final injections, after binding was complete, was used as the heat of dilution and subtracted from each data point. The isotherms were analyzed with Origin Pro data analysis software using a one-site fitting model, which uses a least-square fitting equation. Titrations of DTPA into S33D Zn_7_MT-3 were repeated four times and the reported errors are the standard deviations for the measurements.

## Results

Given the potential importance of the MT-3-actin interaction in explaining the unique biological function of MT-3, we thought it important to characterize how actin polymerization is affected by purified MT-3, without a large GFP tag and in well-defined metalation states.

### Copper ions alter actin filament morphology

Before testing the effect of MT-3 on actin polymerization, we first tested the effect of relevant divalent metal ions in the pyrene-actin polymerization assay. The pyrene assay allows real-time monitoring of the transition from monomeric actin to filamentous actin using the increase in fluorescence of pyrene-labeled actin upon its incorporation into the filament [45,46]. In our experiments, 5% of the actin monomers were pyrene-labeled at residue Cys374, and the remaining 95% were unlabeled. We measured pyrene-actin fluorescence over time in the presence of Zn^2+^ and Cu^2+^, both with and without the actin-binding protein profilin, which is present at high concentration in non-muscle cells and suppresses spontaneous actin filament nucleation (reviewed in [47]). In our experiments, Cu^2+^ addition caused rapid polymerization after a long lag period, an effect that was not observed when Zn^2+^ was added (Fig. 2A). The Cu^2+^-induced increase in the polymerization rate was more prominent in the presence of profilin (Fig. 2B). Our standard polymerization solution contains HEPES buffer, which can be oxidized by Cu^2+^ to form insoluble Cu^+^ species [48–50]. Therefore, we repeated the profilin/actin polymerization assay using MOPS (Fig. S1), a buffer that does not react with Cu^2+^ [49]. The effect of Cu^2+^ on polymerization kinetics was similar in MOPS (Fig. S1) and HEPES (Fig. 2B), indicating that Cu^2+^-induced acceleration of actin polymerization is not an artifact of the unique redox chemistry between Cu^2+^ and HEPES. The effect of Cu^2+^ on actin filaments was suppressed by the addition of 1 mM EGTA in the polymerization buffer (Fig. 2A,B, dashed purple lines).

The pyrene assay is a bulk measurement that provides only the total concentration of monomer in the polymer form but does not provide information about the length distribution of the filaments. We observed a long lag time before the acceleration of actin polymerization over baseline rates, which is consistent with Cu^2+^ ions causing fragmentation of spontaneously-nucleated actin filaments. Fragmentation would create more barbed ends and thus increase the rate of filament growth [51]. To test this model, actin filaments were imaged using TIRF microscopy at two different points in the time course of polymerization (marked by stars in Fig. 2B). The addition of Cu^2+^ ions induced short filaments (Fig. 2C), consistent with electron microscopy of Cu-treated actin filaments [36]. These filaments were notably shorter than those observed in the presence of the formin DIAPH2, one of the human Diaphanous-related formins, which generally accelerate both actin filament nucleation and elongation in the presence of profilin [52]. The effect of Cu^2+^ ions on filament length was reversed by adding EGTA (Fig. 2C). Taken together, the kinetics observed in the pyrene-actin assays (Fig. 2A-B) and the short filament lengths observed by microscopy are consistent with filament fragmentation by Cu^2+^ ions. Spectroscopic data also indicate that Cu^2+^ binding affects intermolecular interactions within the actin filament [53].

### MT-3 reverses the effect of copper ions on actin filament morphology

We next tested whether MT-3 could reverse the effects of Cu^2+^ on actin filament length. In reconstituted systems, MT-3 scavenges Cu^2+^ from pathological protein aggregates such as amyloid *β*-peptides and *α*-synuclein [37,39]. First, we tested whether, under our assay conditions, Zn_7_MT-3 forms a stable complex with actin filaments. We used high-speed co-sedimentation and SDS-PAGE with monobromobimane modification. Under standard, reducing SDS-PAGE conditions, MT-3 runs on the gel anomalously as a diffuse band and is poorly stained by Coomassie, presumably due to its hydrophilicity and lack of aromatic residues. Monobromobimane is a thiol-reactive dye that increases the hydrophobicity of MT-3 and allows detection by Coomassie staining [43]. As expected, actin is present in the pellet fraction after high-speed centrifugation, even in the presence of Cu^2+^, indicating that short filaments are sedimented under these conditions (Fig. 3A). When equimolar actin and Zn_7_MT-3 were mixed, Zn_7_MT-3 was detected in the supernatant fraction in both the presence and absence of Cu^2+^ ions (Fig. 3A; see below for a wider concentration range of Zn_7_MT-3).

To track the binding partner for copper in our co-sedimentation assays, we analyzed the supernatant and pellet fractions using inductively coupled plasma mass spectrometry (ICP-MS). In the absence of Zn_7_MT-3, at least 75% of the detected copper was in the pellet fraction (blue markers in Fig. 3B), consistent with Cu^2+^ bound tightly to actin filaments and therefore co-sedimenting with actin. In the presence of Zn_7_MT-3, less than 2% of the detected copper was in the pellet fraction (blue markers in Fig. 3B), suggesting a transfer of copper from actin filaments to soluble Zn_7_MT-3.

We next asked whether the Cu^2+^-induced fragmentation that we had observed by microscopy (Fig. 2C) could be reversed by the addition of Zn_7_MT-3. In these experiments, F-actin was mixed with Cu^2+^ for 5 minutes before addition of either Zn_7_MT-3 or EGTA. Quantification of actin filament lengths showed that the presence of Zn_7_MT-3 had a modest effect on filament length: 10.0 μm vs 7.8 μm average length without and with Zn_7_MT-3, respectively (Fig. 3C,D). With exposure to only Cu^2+^, short filaments were observed (4.7 μm; Fig. 3C,D). Addition of either Zn_7_MT-3 or EGTA, to the Cu^2+^-fragmented actin resulted in longer actin filaments, presumably through re-annealing [54] following chelation of the Cu^2+^ (Fig. 3C,D). Altogether, the data in Figure 3 are consistent with the transfer of copper ions from F-actin to Zn_7_MT-3 in a mechanism that does not involve a stable MT-3-F-actin complex, similar to other metal swap reactions that have been reported for MT-3 [55].

### Recombinant MT-3 does not bind actin filaments or accelerate actin polymerization

Based on a report that GFP-MT-3 co-sediments with F-actin in astrocyte lysates [27], we were surprised to see minimal co-sedimentation of Zn_7_MT-3 and F-actin (Fig. 3A). To further explore this putative binding interaction, we carried out a co-sedimentation assay using an extended range of Zn_7_MT-3 concentrations (from 0.5:1 to 4:1 Zn_7_MT-3:actin). At all concentrations tested, Zn_7_MT-3 was detected only in the supernatant (Fig. 4). A known actin-bundling protein, fascin, was used as a positive control and was detected in the pellet in the presence of F-actin (Fig. 4). From these data, we conclude that either purified, recombinant Zn_7_MT-3 does not form a stable complex with F-actin, or, if it does form a complex, the equilibrium dissociation constant (K_d_) is much greater than 20 μM and outside the limit of detection for this assay.

**Figure 4.**
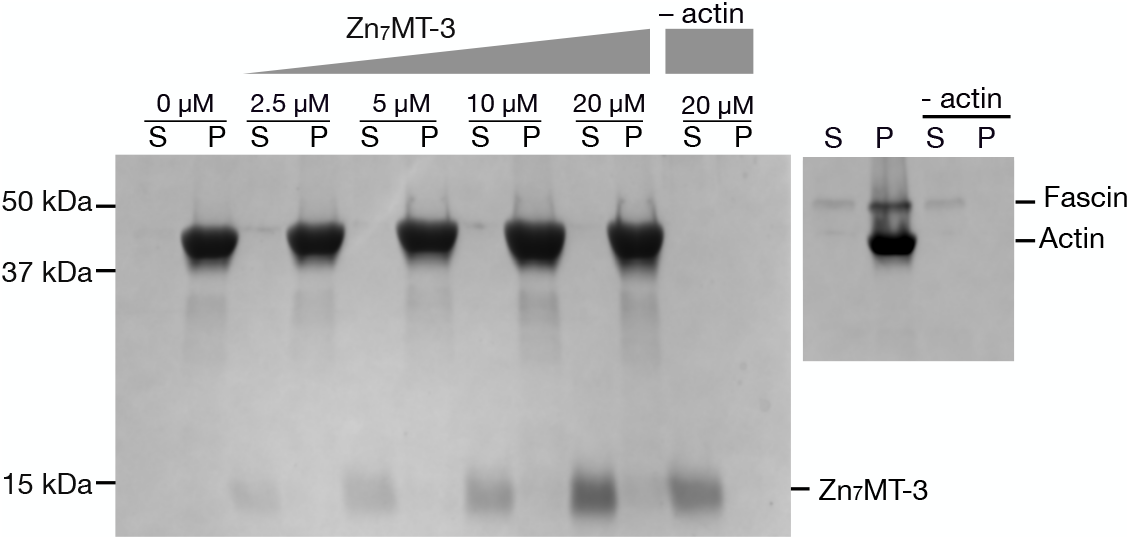
Co-sedimentation assay with Zn_7_MT-3 and F-actin. Zn_7_MT-3 (0, 2.5, 5, 10, or 20 μM) was incubated with 5 μM F-actin and then centrifuged at 90,000 ×*g* for 90 min. The presence of Zn_7_MT-3 in supernatant (S) and actin in pellet (P) implies either (1) that there is not a direct Zn_7_MT-3/F-actin interaction or (2) that the complex has a dissociation constant (K_d_) much greater than 20 μM. Fascin (5 μM) was used as control protein which binds to F-actin (right image). Samples were labeled with monobromobimane, separated by SDS-PAGE, and Coomassie stained. The pellets were concentrated 4-fold compared to the supernatants.

Next, we tested the effect of MT-3 on actin polymerization kinetics using the pyrene-actin polymerization assay, both with and without profilin. For this assay, we prepared MT-3 in three metalation states: Zn_7_MT-3 since this form was reported to interact with actin [27], Cu_4_Zn_4_MT-3 to mimic the physiological form of MT-3 [17,29], and Pb_7_MT-3 since Pb^2+^ can displace Zn^2+^ bound to MT-3 [56] and therefore may be bound to MT-3 *in vivo* following lead exposure. No change in the actin polymerization rate was observed at any concentration of MT-3, in any metalation state (Fig. 5). Representative traces are shown in panels 5A, B, D, E, G, and H, and replicates for each condition are shown in panels 5C, F, and I. As a positive control, we added 5 nM of the formin DIAPH2 to demonstrate that the profilin-actin was competent for polymerization (panels 5B, E and H). These data are different from the report that astrocyte-derived GFP-MT-3, which increased the pyrene-actin signal by ≈30%, indicative of an increase in the steady-state concentration of F-actin [27] (see below for further discussion).

**Figure 5.**
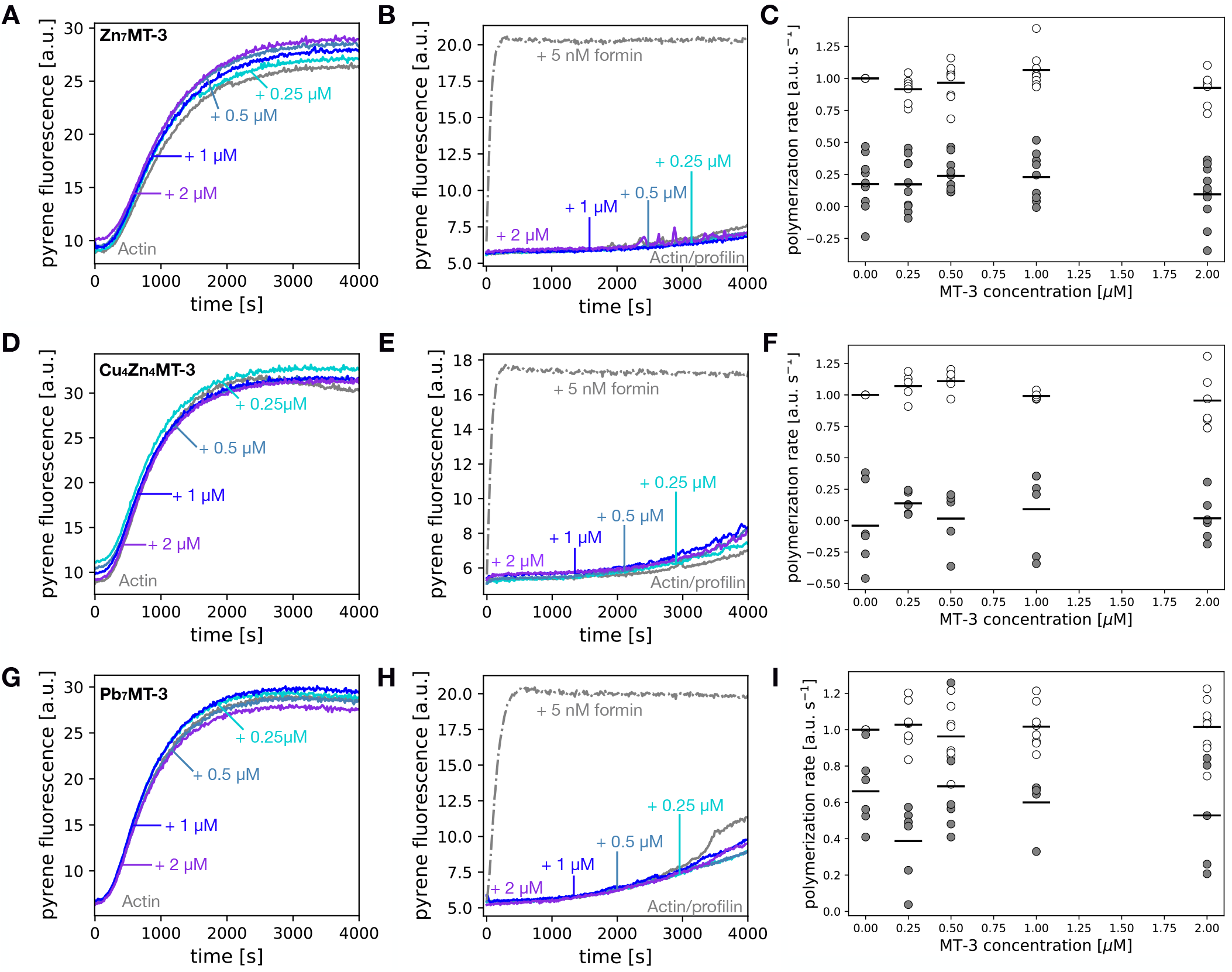
Recombinant MT-3 does not accelerate actin polymerization. Actin polymerization is not affected by the presence of MT-3 in the (A-C) Zn_7_MT-3, (D-F) Cu_4_Zn_4_MT-3, and (G-I) Pb_7_MT-3 forms. Representative raw polymerization traces are shown with varying amounts of MT-3 with either (A, D, F) 4 μM actin or (B, E, G) 3 μM actin/6 μM profilin. For the calculation of polymerization rates (C, F, I), the slope of best fit line at 500 s for 4 μM actin (open circles) was normalized to the actin alone condition for each experiment. The raw slope values at 1000 s for the profilin/actin condition are plotted (filled circles).

The pyrene-actin polymerization assays in Figure 5 were carried out at in the presence of 0.2 mM ATP, which binds MT-2 with a K_d_ of 176 μM and causes large-scale structural changes in MT-2 [57]. We reasoned that ATP in our polymerization assays may affect the MT-3-induced acceleration of actin polymerization, either promoting or inhibiting MT-3-actin interactions. Therefore, we varied the concentration of ATP from 0.01 to 1 mM. Zn_7_MT-3 did not accelerate the rate of actin polymerization at any ATP concentration (Fig. S2). While we cannot rule out an MT-3-ATP interaction, if there is any ATP binding, it does not affect MT-3-actin interactions in the range of concentrations tested (0.01–1 mM).

The difference in actin regulation between our recombinant, bacterially expressed MT-3 (Fig. 5) and astrocyte-derived GFP-MT-3 [27], led us to speculate that recombinant MT-3 was missing a crucial post-translational modification that is important for its interaction with the actin cytoskeleton. To our knowledge, the only reported post-translational modification of metallothionein is phosphorylation of MT-1 at serine 32 (Ser33 in MT-3) [58]. This site is phosphorylated by protein kinase C in response to sub-lethal, pre-conditioning stimuli in cultured neurons. This event has been linked to an increased cytosolic concentration of Zn^2+^ [58], suggesting that this Ser32/33 modification may lower the affinity of metallothionein for Zn^2+^. We created a phospho-mimetic version of MT-3 with the mutation Ser33Asp (S33D). First, we measured the affinity of S33D MT-3 for Zn^2+^ using a well-established isothermal titration calorimetry (ITC) chelation assay [34]. From these experiments, we determined the Zn^2+^ binding free energy, *ΔG°_ITC,S33D_* = −10.1 ± 0.3 kcal·mol^−1^ (Fig. S3), for comparison to that of WT MT-3, *ΔG°_ITC,WT_* = −9.5 ± 0.4 kcal·mol^−1^ [34], which give similar binding constants of *K_ITC,S33D_* = 3 (± 1) × 10^7^ and *K_ITC,WT_* = 1.1 (± 0.9) × 10^7^. Surprisingly, the enthalpic contribution to Zn^2+^ binding (*ΔH°_ITC_*) is ≈2 kcal·mol^−1^ more favorable for WT than for S33D. This difference is compensated for by an entropic contribution to Zn^2+^ binding (−T*ΔS°_ITC_*) at 25 °C that is ≈2.5 kcal·mol^−1^ more favorable for S33D than for WT, leading to similar overall binding affinities (Fig. S3B). In pyrene-actin polymerization assays, S33D Zn_7_MT-3, like WT Zn_7_MT-3, did not alter the kinetics of actin polymerization (Fig. S4D-F). Mutating the two N-terminal prolines that are unique to MT-3 alters the structure and dynamics of the *β* domain and abolishes GIF activity [16]. We therefore tested whether a non-GIF mutant of MT-3 (P7S/P9A Zn_7_MT-3) would have distinct interactions with the cytoskeleton. In pyrene-actin polymerization assays, this variant behaved similar to WT Zn_7_MT-3, with no effect on actin polymerization (Fig. S4A-C).

## Discussion

Actin and MT-3 have been linked in a number of studies [24–28]. These links motivated us to investigate the molecular properties underlying the actin-MT-3 interaction and to probe the role of metal ions in this interaction. Overall, our results suggest that recombinant MT-3 and actin do not bind directly. However, due to the fact that they share a common binding partner in copper ions, MT-3 and actin modulate each other’s properties. This is most evident in our experiments that monitored length distribution of actin filaments in the presence of Cu^2+^, with or without Zn_7_MT-3 (Fig. 2). Electron paramagnetic resonance indicates that copper ions bound to actin are in the +2 oxidation state [33]. While it is clear from our microscopy data that the effect of Cu^2+^ on actin filament morphology is altered by Zn_7_MT-3, we also presume that copper transfer to MT-3 alters the MT-3 structure. This assumption is based on the studies of Meloni, Vašák, and coworkers showing that upon mixing Zn_7_MT-3 and either Cu^2+^ or Cu^2+^-protein complexes, Zn^2+^ is released and MT-3 reduces four Cu^2+^ to Cu^+^, forming two disulfide bonds in the *β* domain in the process [38,59]. The coordination geometry and dynamics of the air-stable Cu^+^_4_CysS_5_ cluster in the *β* domain are distinct from the Zn^2+^_3_CysS_9_,’ cluster present in the *β* domain of Zn_7_MT-3 [59]. Further spectroscopic studies would be necessary to confirm that the reaction products of Zn_7_MT-3 with Cu^2+^-actin are similar to those reported for other metal swap reactions [37,39,40].

Our results do not support a model of MT-3 as an actin assembly factor on its own. This functional assignment was first proposed in a study by Koh and coworkers, who reported that GFP-MT-3 enhances actin polymerization in the pyrene-actin assay [27]. However, there are key differences in the experimental conditions. Most notably, we used recombinant, bacterially expressed MT-3, and in the earlier study, GFP-MT-3 fusion protein was purified by GFP-immuno-affinity chromatography from mouse astrocytes [27]. These different expression systems lead to at least three important caveats that may provide insight about a putative actin-MT-3 interaction.

First, astrocyte-derived MT-3 could have post-translational modifications (PTMs) that are essential for MT-3 function and are missing in bacterially expressed MT-3. We evaluated one possible PTM, phosphorylation at Ser33, that had been reported in a separate study of MT-3 in cortical neurons [58]. Despite WT-like Zn^2+^ affinity (Fig. S3), the phospho-mimetic variant (S33D) showed a similar lack of stimulation of actin assembly, as found with WT MT-3 (Fig. S4D-F). This does not rule out the possibility of other key PTMs in astrocyte-derived GFP-MT-3. PTMs on actin, most notably lysine acetylation [60], can modulate its interactions with binding proteins. However, rabbit skeletal muscle actin was used in this study and in the experiments of Koh and coworkers [27]. Therefore, actin isoform or modification does not explain the differences we observe in MT-3’s stimulation of actin polymerization.

Second, it is possible that GFP-MT-3 from astrocytes co-purifies with an essential binding partner, and together they form a ternary complex with actin. Among the binding partners identified for MT-3 in mass spectrometry screening [25] was <u>d</u>ihydropyrimidase-<u>r</u>elated <u>p</u>rotein 2 (DRP-2; also known as <u>c</u>ollapsin <u>r</u>esponse <u>m</u>ediator <u>p</u>rotein 2, CRMP2). DRP-2 promotes axonal growth cone extension [61], and members of the same family have well-characterized, direct effects on actin [62] and microtubule dynamics [63]. Therefore, DRP-2 is a promising candidate for a cytoskeletal co-regulator that may interact with MT-3 and actin, with the caveat that MT-3 was not identified in a mass spectrometry screen of the DRP-2 interactome [64].

Third, in the earlier study MT-3 was fused to GFP at the N-terminus [27]. The MT-3 used in this study was purified with GFP fused to its N-terminus, and then it was cleaved with TEV protease, leaving an N-terminal Gly-Ser with an unblocked N-terminus. This is distinct from the GFP fusion protein used by Koh and coworkers [27] and also from endogenous MT-3, which is N-acetylated at Met1 [17,65]. Therefore, neither this study nor the study of Koh and coworkers [27] capture the most physiologically relevant N-terminus of MT-3. The fact that the N-termini are different in two experiments could point to a role for that region in MT-3-actin interactions, a role supported by data from Koh and co-workers, who show that an N-terminal peptide containing the unique ‘TCPCPT’ in MT-3 is sufficient to block c-Abl/ Factin interactions [27].

In addition to a role for MT-3 in astrocyte actin remodeling, MT-3 is implicated in cytoskeletal regulation during the doming of kidney proximal tubule cells [26,66]. The doming process in cultured cells is a model for vectorial active transport. Garrett and coworkers over-expressed V5-tagged MT-3 in HK-2 proximal tubule cells and were able to co-immunoprecipitate *β*-actin, myosin-9, and enolase 1. Furthermore, MT-3 co-localized with F-actin in the doming HK-2 cell layer [26], and cell lines that did not express MT-3 fail to dome [6,66]. Based on these findings and molecular mechanics docking calculations, Garrett and coworkers proposed a structural model where MT-3 binds to the target-binding cleft of actin between subdomains 1 and 3 via sidechain contacts in the *α* domain of MT-3. Generally, proteins that bind at this site on actin affect polymerization dynamics, either through filament nucleation, monomer sequestration, barbed end capping, or filament severing [67]. However, none of these activities is observed in our assays. Suppression of actin polymerization due to monomer sequestration or barbed end capping could be observed under conditions in which substantial spontaneous filament nucleation occurs (4 μM actin; Fig. 5A, D, G) but is not seen in the presence of MT-3. Filament nucleation or severing would increase the rate of actin polymerization in a pyrene assay. However, acceleration by MT-3 is not observed in the absence (Fig. 5A, D, G) or presence (Fig. 5B, E, H) of profilin.

In summary, purified, recombinant MT-3 does not bind to muscle actin. The Zn_7_MT-3 form of MT-3 removes Cu^2+^ ions from actin filaments, reversing the effects of copper-induced F-actin fragmentation. Previous reports of Zn_7_MT-3-mediated Cu^2+^ scavenging involve proteins such as amyloid-*β*, *α*-synuclein, and prion protein [37,39,40] that are found in the extracellular compartment, a context where the +2 oxidation state of copper may be more prevalent than in the reducing environment of the cytoplasm, where actin is found [68]. Nonetheless, there are particularly high concentrations of Cu ions in the central nervous system [69], and in cultured neurons, where Cu (of unknown oxidation state) is concentrated at the necks of dendritic spines, a particularly actin-rich structure [35]. We posit that understanding the interplay of Cu, MT-3, and the actin cytoskeleton is important to understand MT-3’s role as a GIF and generally in the CNS and kidney in health and disease.

## Acknowledgements

The authors are grateful to Naomi Courtemanche for sharing the fascin plasmid and to Jamie Ross and Benjamin Bostick for assistance with ICP-MS. RL, RNA, and CLV were supported by NSF grant CHE-1710176. MRM and DEW were supported by NSF grant CHE-1904705. The microscope was purchased with funds from NSF grant DBI-1828264.

## Accession IDs

For recombinant proteins used in this study, the mouse MT-3 amino acid sequence corresponded to UniProt P28184, *S. pombe* profilin to UniProt P39825, and human fascin1 to UniProt Q16658, human DIAPH2 to UniProt O60879-2 (residues 553–1096).

---

**Figure S1.**
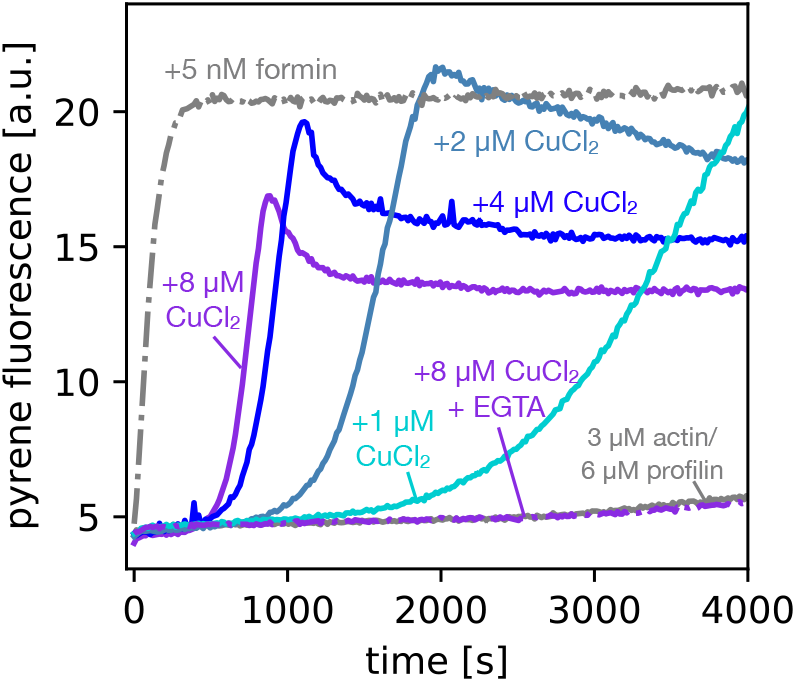
Effect of CuCl_2_ on actin polymerization in the presence of MOPS buffer. Conditions were identical to Figure 2B (3 μM actin, 6 μM profilin), with the exception that 10 mM MOPS was used instead of 10 mM HEPES in the polymerization buffer.

**Figure S2.**
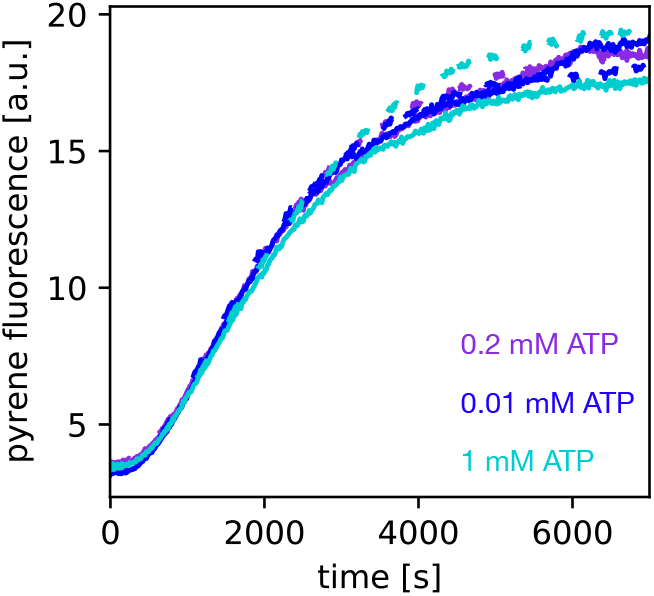
Varying ATP concentration does not alter the effect of Zn_7_MT-3 on actin polymerization. Actin polymerization was measured in the presence of varying ATP concentration (0.01 mM ATP, 0.2 mM ATP, and 1 mM ATP). For each ATP concentration, the dashed and solid line represents the assay with and without 2 *μ*M Zn_7_MT-3, respectively. For all experiments, the actin concentration was 2 *μ*M (5% pyrene-labeled).

**Figure S3.**
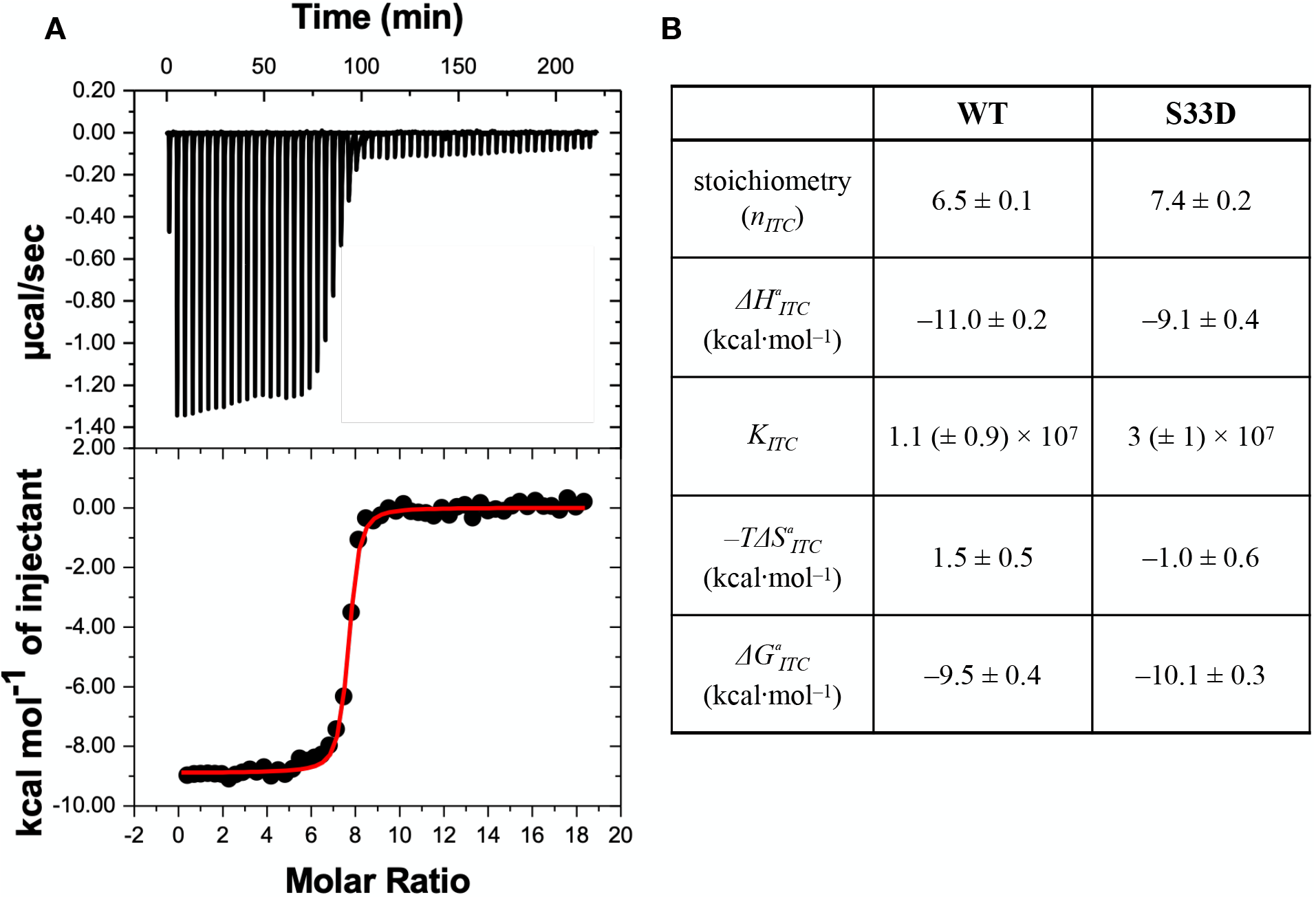
Thermodynamics of Zn^2+^ binding to S33D MT-3 from Zn^2+^ chelation ITC data. (A) Representative ITC thermogram for titration of the chelator DTPA into S33D Zn_7_MT-3 at 25°C; 700 *μ*M DTPA and 5 *μ*M S33D Zn_7_MT-3, both in 100 mM BisTris, 150 mM NaCl, pH 7.4; (B) Average thermodynamic parameters for Zn^2+^ binding to S33D MT-3 from the analysis of ITC data from four replicate chelation titrations and comparison to average thermodynamic parameters for Zn^2+^ binding to WT Zn_7_MT-3 [34].

**Figure S4.**
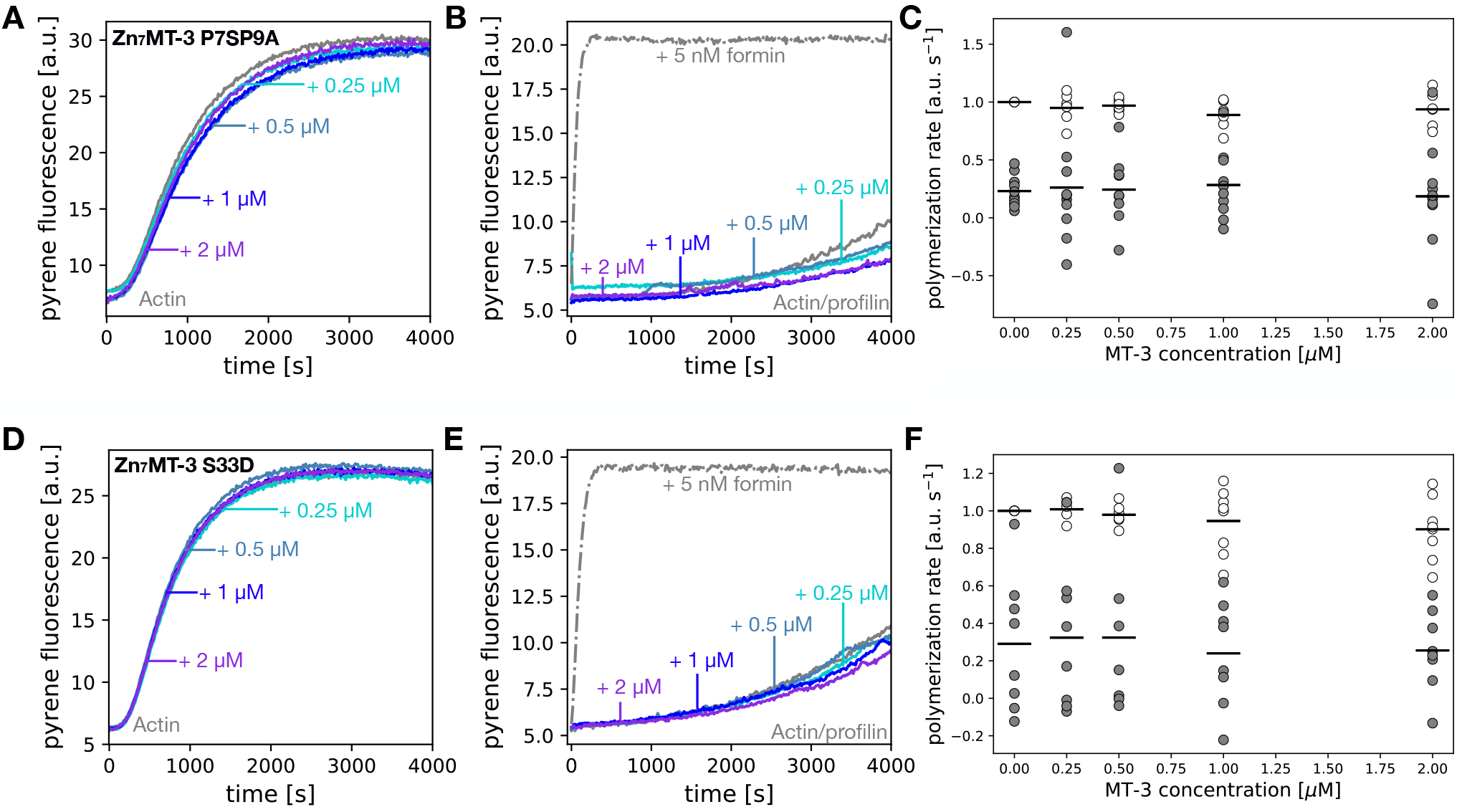
The non-GIF mutant MT-3 (P7S/P9A) and phosphomimetic mutant MT-3 (S33D) do not affect actin polymerization. Actin polymerization is not affected by Zn bound non-GIF mutant MT-3 (Pro7Ser/ Pro9Ala) (A-C) and phosphomimetic mutant MT-3 (Ser33Asp) (D-F) similar to the WT Zn_7_MT-3 (Figure 5). Representative raw polymerization traces are shown with varying amounts of MT-3 with either (A, D) 4 μM actin or (B, E) 3 μM actin/6 μM profilin. For the calculation of polymerization rates (C, F), the slope of best fit line at 500 s for 4 μM actin was normalized to the actin alone condition for each experiment. The raw slope values at 1000 s for the profilin/actin condition are plotted.

## SI Text file: sequences of MT-3 proteins used in this study

Codons for His_6_-GFP-tev-**MT-3** (based on accession *#* NP_038631.1):

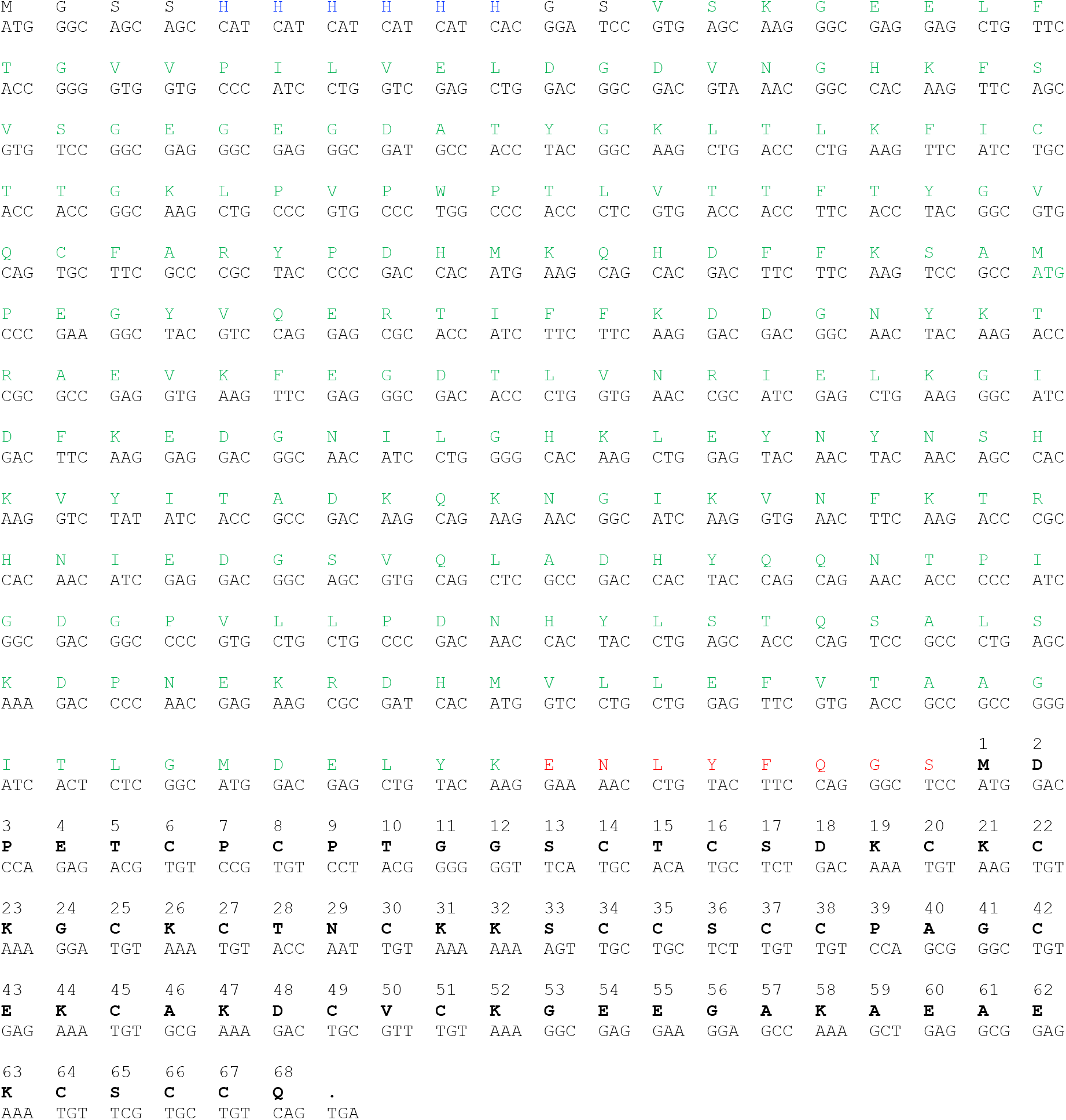

Cleaved WT MT-3 used in experiments:

~~~
GSMDPETCPCPTGGSCTCSDKCKCKGCKCTNCKKSCCSCCPAGCEKCAKDCVCKGEEGAKAEAEKCSCCQ
~~~

Codons for His_6_-GFP-tev-**MT-3**-S33D :

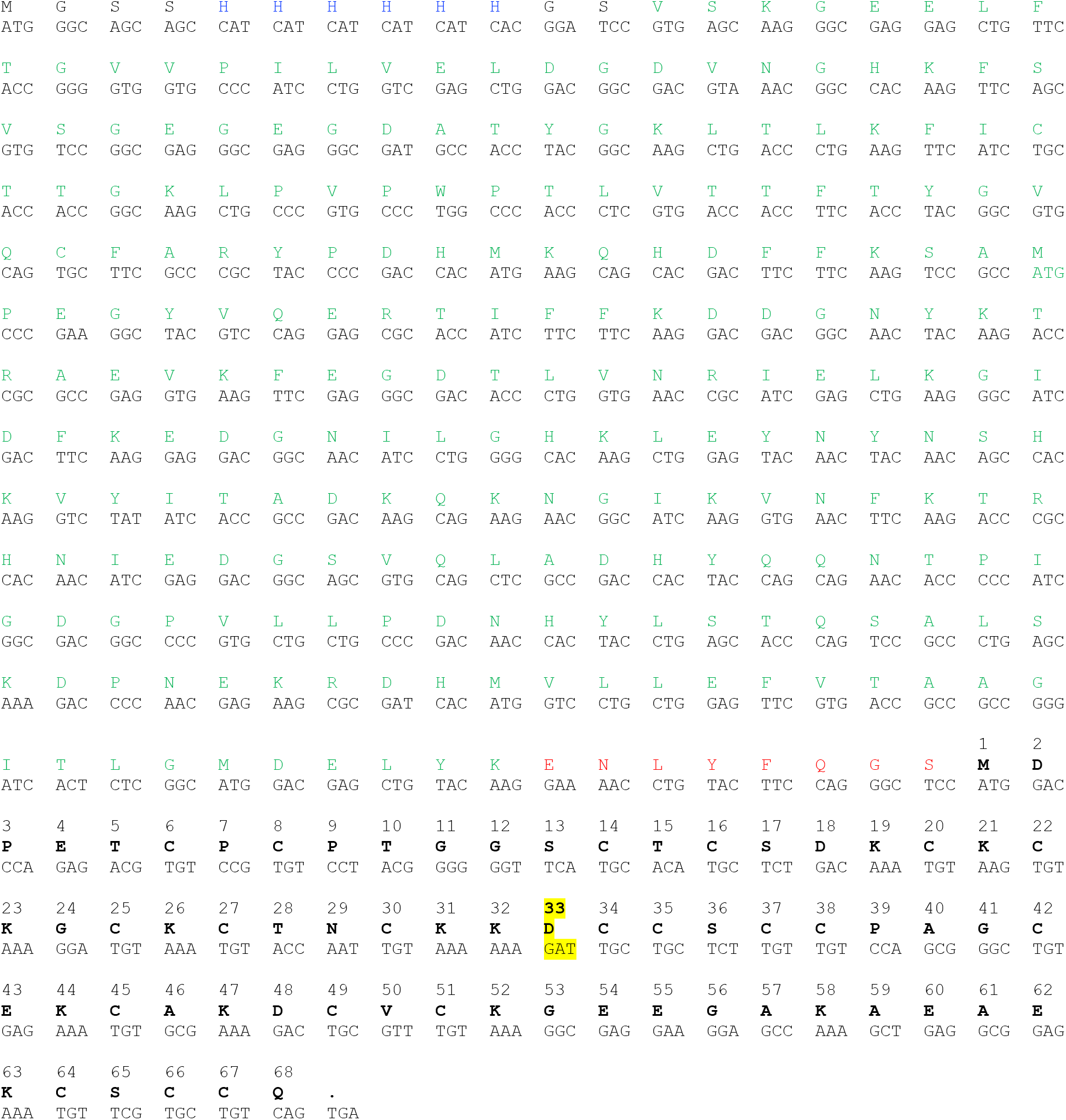

Cleaved S33D MT-3 used in experiments:

~~~
GSMDPETCPCPTGGSCTCSDKCKCKGCKCTNCKKDCCSCCPAGCEKCAKDCVCKGEEGAKAEAEKCSCCQ
~~~

Codons for His_6_-GFP-tev-**MT-3**-P7S/P9A :

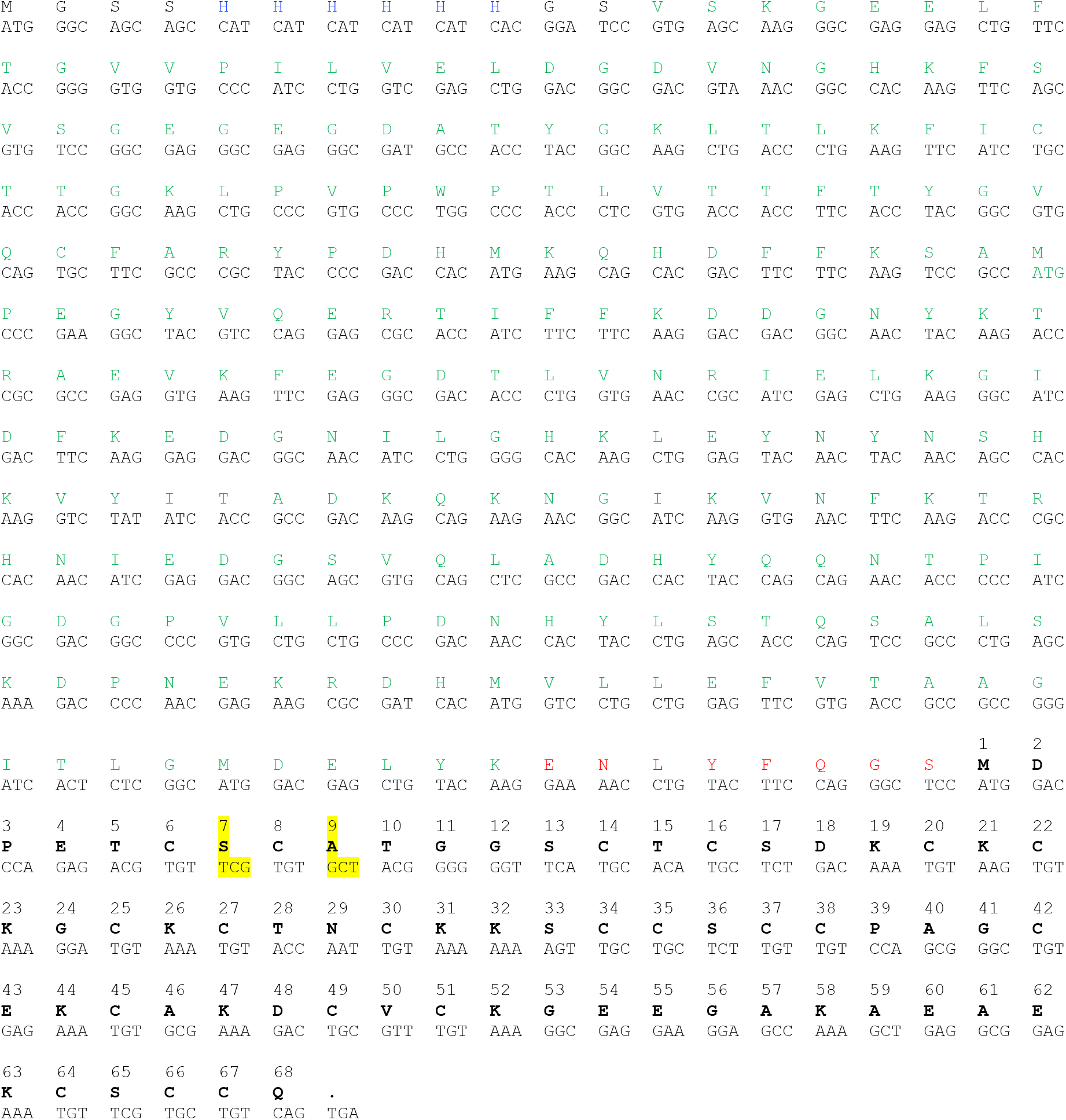

Cleaved P7S/P9A MT-3 used in experiments:

~~~
GSMDPETCSCATGGSCTCSDKCKCKGCKCTNCKKSCCSCCPAGCEKCAKDCVCKGEEGAKAEAEKCSCCQ
~~~

